# Ghrelin enhances food odor conditioning in healthy humans: an fMRI study

**DOI:** 10.1101/311738

**Authors:** Jung Eun Han, Johannes Frasnelli, Yashar Zeighami, Kevin Larcher, Julie Boyle, Ted McConnell, Saima Malik, Marilyn Jones-Gotman, Alain Dagher

## Abstract

Vulnerability to obesity includes eating in response to food cues, which acquire incentive value through conditioning. The conditioning process is largely subserved by dopamine, theorized to encode the discrepancy between expected and actual rewards, known as the reward prediction error (RPE). Ghrelin is a gut-derived homeostatic hormone that triggers hunger and eating. Despite extensive evidence that ghrelin stimulates dopamine, it remains unknown in humans if ghrelin modulates food cue learning. Here we show using functional magnetic resonance imaging that intravenously administered ghrelin increased RPE-related activity in dopamine-responsive areas during food odor conditioning in healthy volunteers. Participants responded faster to food odor-associated cues and perceived them to be more pleasant following ghrelin injection. Ghrelin also increased functional connectivity between hippocampus and ventral striatum. Our work demonstrates that ghrelin promotes the ability of cues to acquire incentive salience, and has implications for the development of vulnerability to obesity.

## Introduction

Accumulating evidence from psychology, cognitive neuroscience, genetics, and neuroimaging has established the role of higher-level cognitive and emotional brain systems in the maintenance of energy balance in humans. Homeostatic peptides from the periphery convey energy balance information to the brain. In order for this information to affect food intake it must influence brain circuitry involved in decision-making and motivation.

Exposure to cues associated with palatable food can evoke motivation to eat, and eventually lead to weight gain (Boswell and Kober, 2016; Jansen et al., 2016; Petrovich et al., 2007). The cue-potentiated feeding response results from conditioning whereby neutral cues acquire incentive value after being repeatedly paired with ingestion of food. Such cues include the sight, smell and flavour of food. The ability of food cues to become conditioned as well as their subsequent potency to elicit feeding is greater in the hungry state (Balleine, 1992; LaBar et al., 2001). A likely candidate mediating the interaction of hunger and food cue conditioning is the hormone ghrelin.

Ghrelin is a stomach-derived peptide hormone that elicits hunger and feeding by acting on the brain (Müller et al., 2015). It binds to a unique receptor, the growth hormone secretagogue receptor (GHSR), expressed densely in brain areas involved in feeding and energy balance, such as the hypothalamus and nucleus of the solitary tract (Mason et al., 2014). Ghrelin levels rise prior to scheduled mealtimes and after fasting, and fall postprandially (Cummings, 2004; Cummings et al., 2001; Muller et al., 2002). Moreover, administration of ghrelin induces hunger and food consumption in the short term while promoting fat accumulation in the long term (Druce et al., 2005; Nakazato et al., 2001; Tschöp et al., 2000; Wren et al., 2001). Ghrelin signals several different types of information that affect the motivation to eat, notably the immediate availability of food, the timing of an expected meal, and both short and long-term energy balance status (Müller et al., 2015). There is much evidence that ghrelin acts not only on the homeostatic hypothalamic-brainstem circuits that regulate energy balance but also on systems involved in learning and motivation, notably the ventral tegmental area (VTA), striatum and hippocampus, to influence food cue reactivity. More specifically, ghrelin may increase the motivational salience of food cues by stimulating dopaminergic neurons in the VTA where GHSR are also found (Mason et al., 2014; Perello and Dickson, 2015). Ghrelin injection into the VTA increases activity of dopamine (DA) neurons and triggers DA release in the nucleus accumbens while motivating animals to work harder to obtain food rewards (Abizaid et al., 2006; Jerlhag et al., 2007; Skibicka et al., 2011, 2013). On the other hand, administration of a ghrelin or DA antagonist abolishes the ghrelin-induced increase in food motivated behaviour (Skibicka et al., 2011, 2013). These findings from animal studies are corroborated by fMRI studies in humans. High levels of ghrelin in healthy volunteers, as a result of fasting or intravenous ghrelin injection, appear to enhance the incentive salience of food cues, as reflected by stronger activity in response to food images in brain regions such as the orbitofrontal cortex (OFC), striatum and hippocampus, and greater subsequent recall of the food images (Goldstone et al., 2014; Kroemer et al., 2013; Malik et al., 2008).

Ghrelin’s ability to stimulate DA has implications not only for its influence on responses to learned cues associated with food but also for the food cue conditioning process. Associative learning is theorized to be driven by the discrepancy between the expected value assigned to the cue and the value of the actual reward outcome, known as the reward prediction error (RPE) (Glimcher, 2011; Schultz, 2016). Phasic firing of DA neurons in the VTA is thought to encode the RPE, through which the DA system contributes to acquisition and update of reward-cue associations. DA phasic signaling in response to food cues is augmented by central ghrelin injection (Cone et al., 2015). GHSR knockout mice, on the other hand, do not demonstrate release of accumbens DA upon exposure to food (Egecioglu et al., 2010). These findings collectively suggest that ghrelin may promote food-cue associative learning by enhancing the phasic RPE signal.

Another region implicated in food-cue related associative learning is the hippocampus (Kanoski and Grill, 2017). There is also a high concentration of GHSR in the hippocampus, where ghrelin can increase spine density and improve learning and memory, possibly by modulating dopamine signaling (Diano et al., 2006; Li et al., 2013; Ribeiro et al., 2014). Conditioned feeding, which occurs in response to learned food-cue associations, is increased in rats upon ghrelin injection into the ventral hippocampus (Hsu et al., 2016; Kanoski et al., 2013). To date, the influence of ghrelin on food-related conditioning has only been tested in animals. Ghrelin injection enabled conditioned place preference to high fat food in wild-type mice but not in GHSR knockouts (Perello et al., 2010). Moreover, caloric restriction associated with high levels of endogenous ghrelin failed to induce conditioned place preference in GHSR-null mice or those treated with a GHSR-antagonist during the conditioning phase.

Whether ghrelin also modulates food-cue associative learning in humans remains unexplored. In the midst of an escalating global obesity epidemic, this is an important question to address given the role of excessive food cue learning and reactivity observed in obese individuals (Boswell and Kober, 2016; Jansen et al., 2016) and evidence of impaired ghrelin signaling in obesity (Zigman et al., 2016). Here we test the ability of the orexigenic peptide ghrelin to promote Pavlovian conditioning to food odors by increasing neural reward prediction error activity in dopaminergic projection sites such as ventral striatum (VStr) and hippocampus. This work attempts to make a link between homeostatic signaling and learning systems that help shape food behavior.

Following intravenous administration of ghrelin (1μg/kg) or saline on two separate days, thirty-eight subjects underwent functional magnetic resonance imaging (fMRI) while they learned to associate neutral abstract images with food or non-food odors. Participants rated pleasantness of the images throughout the scan and again 24 hours after each scan session. It was hypothesized that ghrelin would enhance conditioning of cues paired with food, but not non-food, odor via an effect on dopaminergic brain regions.

## Results and Discussion

### Ghrelin Increases Subjective Hunger and Elevates Growth Hormone and Cortisol

Subjective hunger ratings were collected using a visual analogue scale (VAS) throughout the experiment. We observed significant main effects of condition and time (F(1,33)=11.32, p<0.01 and F(3,99)=108.88, p<0.001 respectively) as well as a significant interaction between the two factors (F(3,99)=4.26, p<0.01; see Figure S1(A)). Participants were least hungry after eating breakfast, which was provided 3 hours before ghrelin or saline administration (ps<0.001). Their pre-scan (post-injection) hunger ratings were also lower compared to pre-breakfast and post-scan ratings (ps<0.001). Consistent with the role of ghrelin, post-injection and post-scan hunger ratings were higher in the ghrelin versus saline condition (t(33)=4.83, p<0.001 and t(33)=2.16, p<0.05 respectively). VAS ratings of boredom and irritability did not differ between conditions (Figure S1 (B) & (C)).

Given that ghrelin binding to central nervous system GHSR triggers growth hormone (GH) secretion (Arvat et al., 2001), another way to measure a brain effect of ghrelin is to examine associated changes in GH levels. Blood samples were withdrawn before ghrelin injection (before the MRI scan) and after the scan to quantify levels of GH. As illustrated in Figure S2(A), we observed significant main effects of condition (F(1,25)=31.90, p<0.001) and time (F(1,25)=34.26, p<0.001) and a significant interaction between the two variables (F(1,25)=35.38, p<0.001). As expected, post-scan growth hormone levels were significantly higher following ghrelin compared to saline administration (t(25)=5.91, p<0.001). One participant did not show the expected growth hormone response to ghrelin injection (pre-scan: 3ug/L, post-scan: 2.12ug/L) and was excluded from further analyses.

In line with previous findings (Schmid et al., 2005; Takaya et al., 2000), ghrelin also increased levels of salivary cortisol (Figure S3). We observed significant main effects of condition and time (F(1,32)=12.63, p<0.01 and F(2,64)=3.50, p<0.05 respectively) as well as a significant interaction between the two variables (F(1.32, 42.22)=20.08, p<0.001). At post-scan, cortisol levels were significantly greater following ghrelin than saline infusion (t(32)=5.88, p<0.001).

### Ghrelin Reduces Response Time to Food Odor-Paired Cues and Intensifies Their Pleasantness

We administered a food odor conditioning task during fMRI (Figure 1B) on two different days, following ghrelin or saline intravenous injection (single-blind and counterbalanced). During the task, participants were presented with a series of trials in which one of four abstract images was followed, 50% of the time, by one of two food or two non-food odors (with odorless air being delivered in the remaining trials), or one of two abstract images that invariably cued delivery of odorless air. In all, there were six images and four odors. Participants were instructed to indicate using a MRI-compatible mouse-like device whether each image was composed of straight or curvy lines. This allowed us to examine reaction time, frequently used as an index of learning during classical conditioning. As illustrated in Figure 2A, z-transformed reaction time decreased over the course of the task, regardless of odor type and condition (F(2.74, 76.72)=6.63, p<0.005). We also observed significant interactions between time and odor type (F(12, 336)=3.35, p<0.001) and between condition and odor type (F(2, 56)=3.48), p<0.05; see Figure 2B). Post-hoc paired t tests revealed that the difference in response time between the food and non-food trials differed significantly between the ghrelin and saline conditions (t(28)=-2.47, p<0.05): following ghrelin infusion, subjects responded faster toward food-related images compared to those paired with non-food odors (t(28)=-1.87, p=0.07) while in the saline condition the response time was (not significantly) lower for the non-food odor-paired images (t(28)=1.70, p=0.1). Furthermore, the reaction time difference between food and air trials differentiated the ghrelin and saline conditions (t(28)=-2.20, p<0.05) such that ghrelin induced faster reaction time on the food compared to air trials (t(28)=-2.63, p<0.05) while no such difference was observed following saline administration (t(28)=0.91, p=0.37).

**Figure 1.**
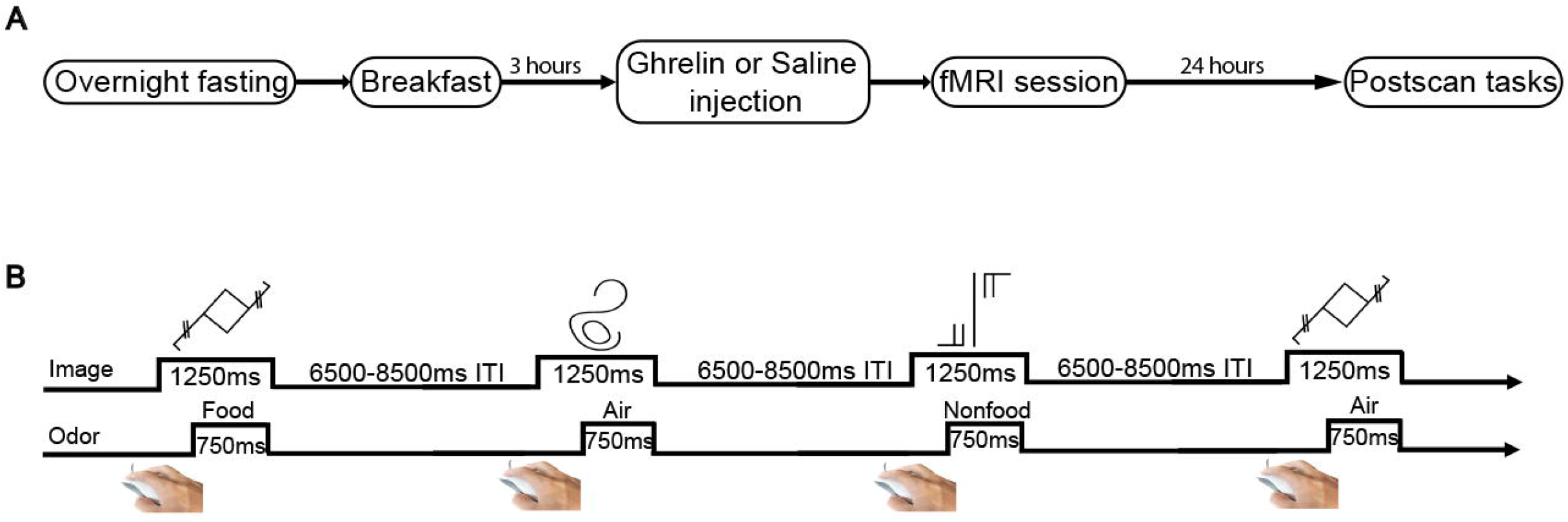
Overview of the protocol. (A) Subjects underwent two sessions, ghrelin or saline, counterbalanced and single-blind, at least 7 days apart. The ghrelin or saline administration and the subsequent fMRI session took place 3 hour post-breakfast. (B) The fMRI conditioning task began with the presentation of an abstract image followed by its corresponding odor or air, ending with an inter-trial blank screen. There were 7 fMRI runs, each of which consisted of 36 image-odor/air trials.

**Figure 2.**
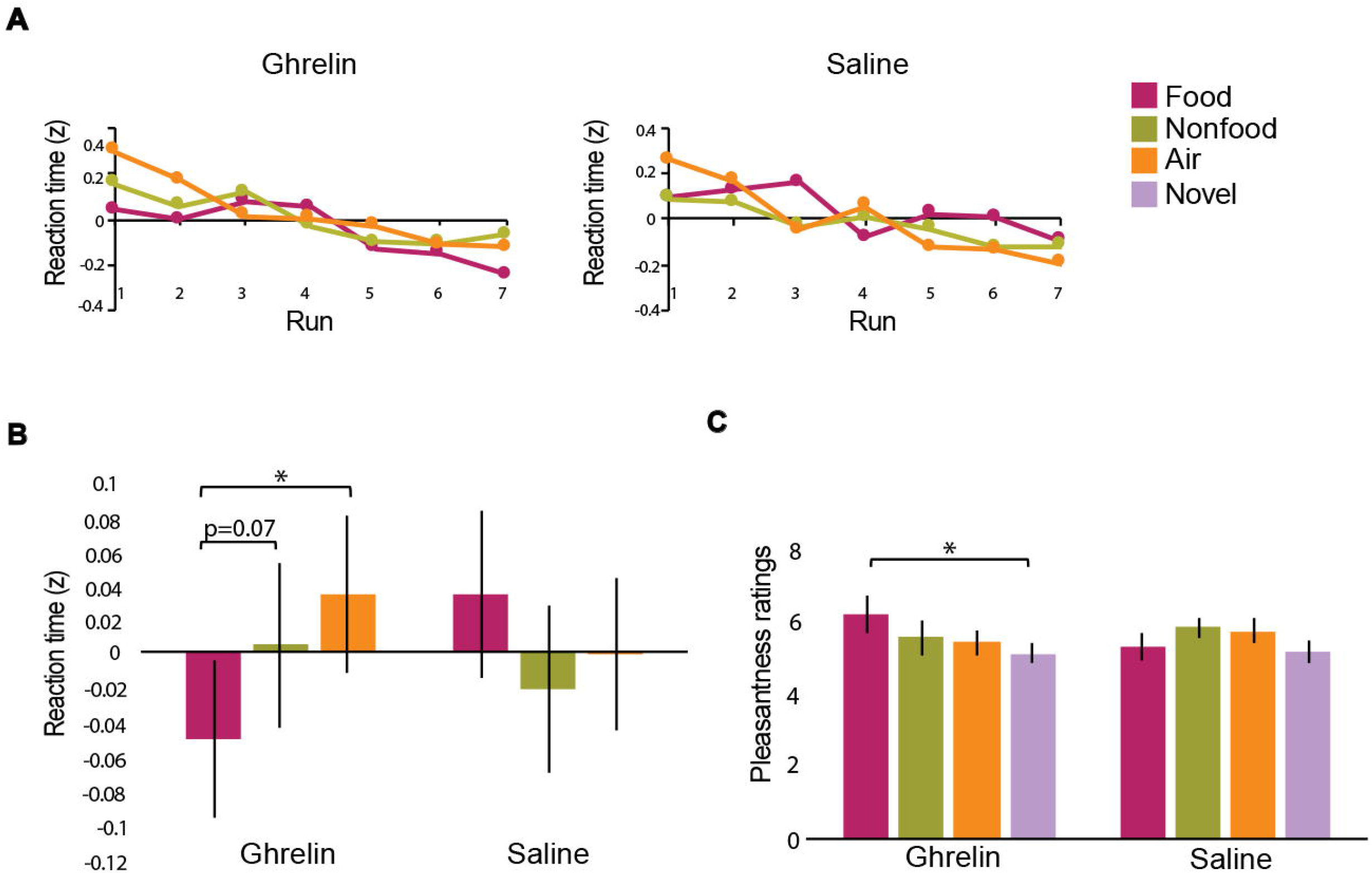
Ghrelin reduces response time to food odor-paired cues and intensifies their pleasantness. (A) In both ghrelin and saline conditions, participants’ reaction times in response to abstract images decreased over the course of the task (F(2.74, 76.72)=6.63, p<0.005). (B) The difference in reaction time between food and non-food trials and that between food and air trials significantly differed between ghrelin and saline conditions (t(28)=-2.47, p<0.05; t(28)=-2.20, p<0.05). Following ghrelin administration, participants responded faster toward food-related images compared to those paired with non-food odors ((t(28)=-1.87, p=0.07) and air (t(28)=-2.63, p<0.05). On the other hand, in the saline condition, food-associated images induced greater response time compared to non-food odor-paired images (t(28)=1.70, p=0.1). (C) The only significant result observed on the hedonic rating task administered after a 24-hour delay was greater pleasantness ratings for food odor-associated images compared to novel ones following the ghrelin session (t(17)=2.14, p<0.05).

We also used hedonic ratings of the abstract images to measure conditioning. Repeated measures ANOVAs conducted on the hedonic ratings collected during the scans and 24 hours after each scan did not yield any significant results. However, paired t tests revealed that after a 24-hour delay, abstract images associated with food odors following ghrelin administration were perceived to be more pleasant than novel images (t(17)=2.14, p<0.05; see Figure 2C). The effect was not significantly different between ghrelin and saline conditions. Taken collectively, faster reaction times and increased liking toward food-associated cues following ghrelin administration suggest that the hormone may accelerate conditioning to food-related stimuli.

### Ghrelin Increases RPE-Associated Activity During Food Odor Conditioning

To induce RPE and reward learning, the fMRI task implemented a 50% reinforcement schedule. In order to map brain activity related to RPE, a group learning rate was first estimated by fitting a Rescorla-Wagner learning model to participant reaction times. We then used the derived learning rate and the model to calculate the trial-by-trial RPE, which was subsequently regressed with brain activation (O’Doherty et al., 2007). In each of the ghrelin and saline conditions (analyzed independently), RPE was positively correlated with activity in a large number of regions including the piriform cortex, amygdala, VStr, putamen, globus pallidus, insula, substantia nigra/VTA, OFC, and anterior and posterior cingulate cortex (Figure 3A, Table 1). In testing the effect of ghrelin on RPE-related activity, we limited the analysis to a group of brain regions previously identified by metaanalysis to subserve RPE, which includes the VStr (Chase et al., 2015). We observed a significant interaction between condition and event type in the region of interest defined by the meta-analysis (F(1,28)=4.60, p<0.05; see Figures 3B & 3C). More specifically, RPE-associated activity following ghrelin injection was greater on food compared to non-food trials (t(28)=2.41, p<0.05). Such difference was absent in the saline condition (t(28)=-0.55, p=0.59). Furthermore, we observed, only in the ghrelin condition, a positive correlation between RPE-associated activity during food-related learning and in-scanner pleasantness ratings of the images associated with food odors (r=0.61; n=14; Figure 3D). An additional analysis demonstrated that condition significantly moderated the relationship between the pleasantness ratings and RPE-related activity on food trials (B: 24.93, *β*: 0.82, p<0.01). Finally, in a separate analysis focusing on the hippocampus, we observed stronger RPE-related activity in the right hippocampus during food-odor conditioning following ghrelin compared to saline administration (p=0.015, FWE after small volume correction using the anatomical hippocampus mask).

**Figure 3.**
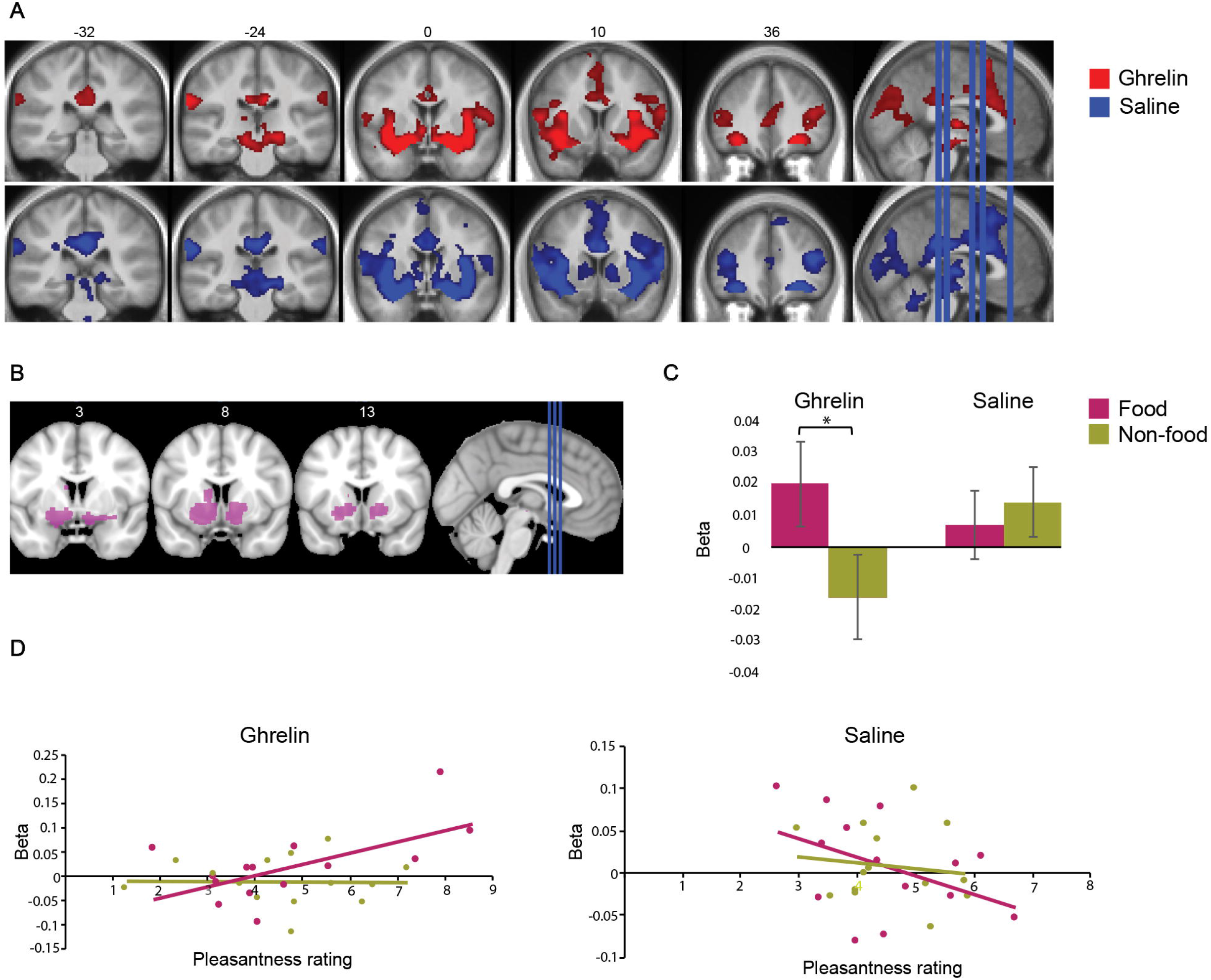
Ghrelin increases RPE-associated activity during food odor conditioning. (A) The whole brain analysis revealed RPE-related activity in a large number of brain regions including the piriform cortex, VStr, putamen, OFC and substantia nigra/VTA in both ghrelin and saline conditions (FDR corrected p<0.05). (B) A mask of the brain regions that were previously identified by a meta-analysis to subserve RPE. We conducted an ROI analysis using the mask and compared RPE-related activity between ghrelin and saline conditions. (C) The ROI analysis revealed that only in the ghrelin condition, RPE-related activity was stronger on food trials compared to non-food trials (t(28)=2.41, p<0.05). (D) In-scanner pleasantness ratings of the abstract images correlated positively with RPE-related activity on food trials following ghrelin infusion (r=0.61). No other significant correlations were revealed.

**Table 1.**
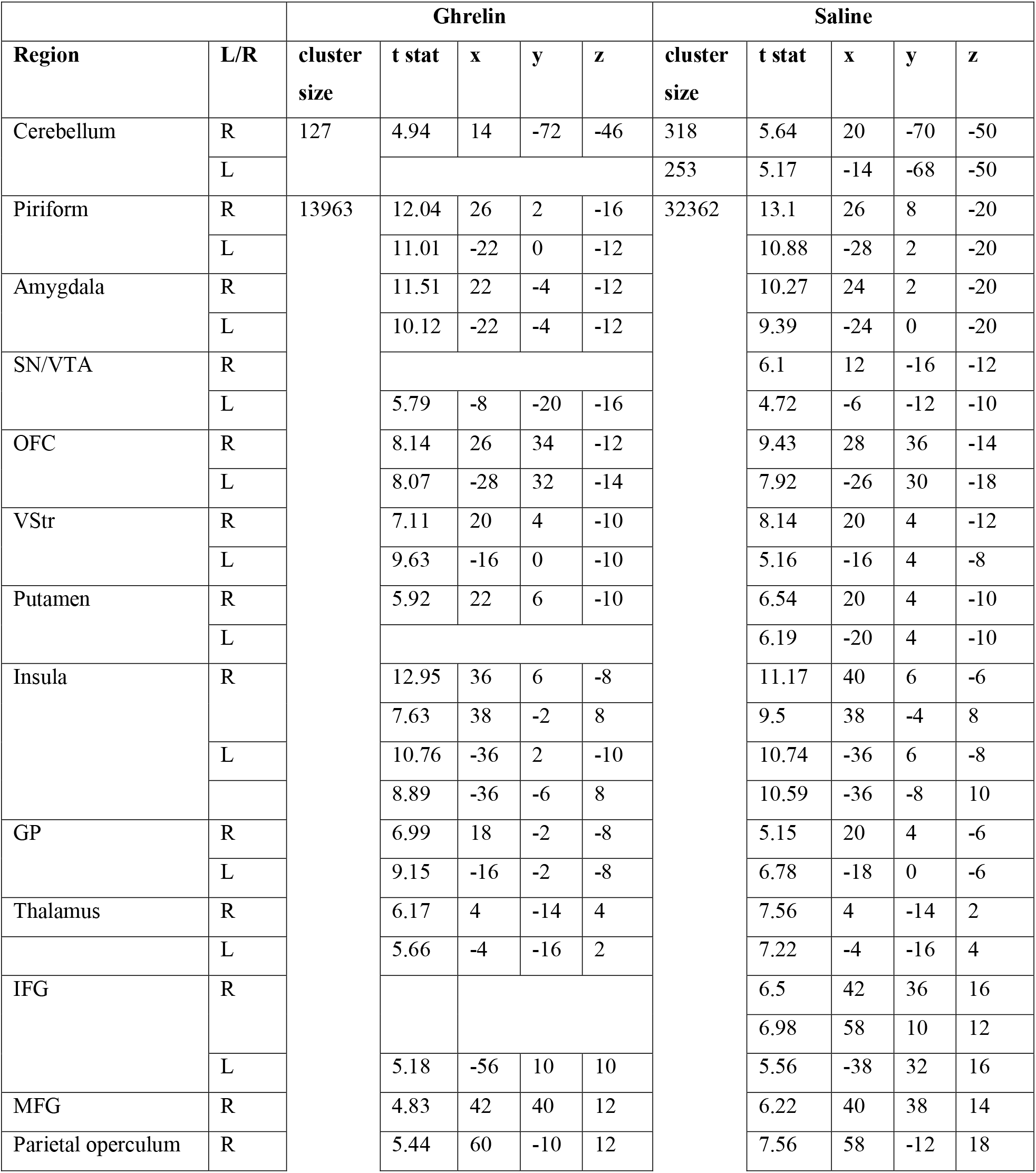
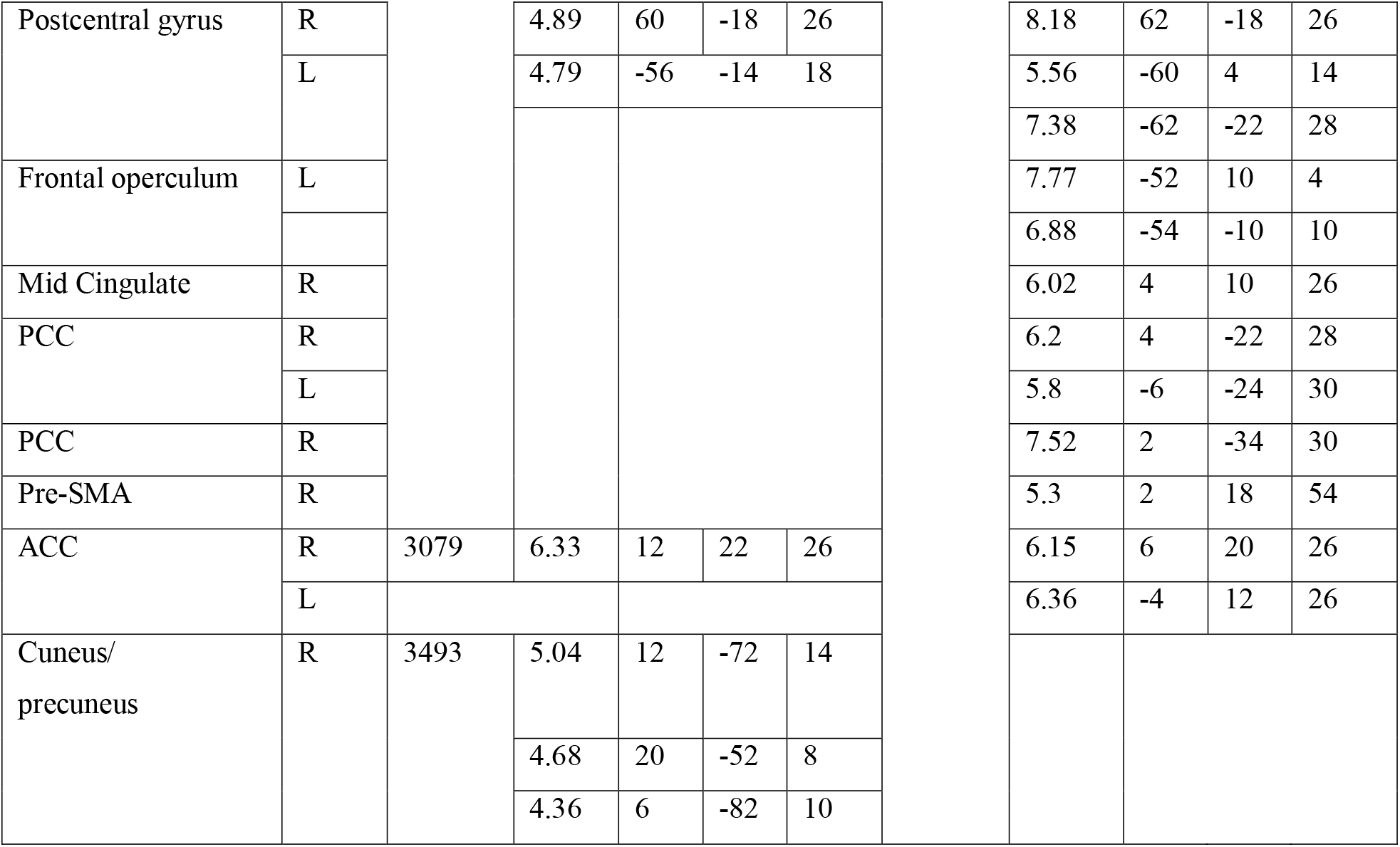
Reward Prediction Error-related activity. SN/VTA: substantia nigra pars compacta / ventral tegmental area, GP: globus pallidus, OFC: orbitofrontal cortex, MFG: middle frontal gyrus, IFG: inferior frontal gyrus, VStr: ventral striatum, PCC: posterior cingulate cortex, Pre-SMA: presupplementary motor area, ACC: anterior cingulate cortex.

Cue-reward associations are thought to be shaped by DA-generated RPE signals (Glimcher, 2011; Schultz, 2016). In human fMRI studies, RPE signals are related to activity in DA-sensitive brain regions such as the striatum, and typically reflect learning (Schonberg et al., 2007). Ghrelin binds to GHSR expressed in the VTA, where it can stimulate DA signaling to promote food cue conditioning (Mason et al., 2014; Perello and Dickson, 2015). Considerable evidence suggests that phasic DA encodes the RPE (Chang et al., 2016; Schultz, 2016). Ghrelin injection is shown to increase phasic DA signaling in response to food cues and to heighten activity in DA-responsive brain regions in humans (Cone et al., 2015; Goldstone et al., 2014; Malik et al., 2008). Furthermore, flavour-nutrient conditioning, a process that mostly implicates olfaction, necessitates D1 receptor-dependent phasic dopamine signaling (Sclafani et al., 2011). Our neuroimaging results extend these findings and provide more direct evidence that ghrelin enhances activity associated with prediction errors for food reward in dopaminergic projection sites, while also accelerating food cue-related learning.

### Ghrelin Heightens the Brain Response Associated with Expected Value Assigned to Food Cues

Successful associative learning is also reflected in the degree to which cues acquire the incentive salience of their associated reward. The reinforcement learning model provides an estimate of expected value assigned to conditioned stimuli (CS) on each trial, hereafter referred to as “CS Value”. The trial-by-trial CS Values were regressed onto fMRI responses, providing another measure of learning-related brain activity. As illustrated in Figure 4A, both conditions induced CS Value-related activity during exposure to the visual cues in a broad range of brain regions including the piriform cortex, insula, globus pallidus, anterior and posterior cingulate cortex and OFC (Table 2). The analysis testing ghrelin’s effects was limited to the two regions of interest previously shown to encode subjective value by meta-analysis, namely the vmPFC and VStr (Bartra et al., 2013). As seen in Figure 4B, CS Value-related activity in the vmPFC was only significant on food-odor trials in the ghrelin condition (t(28)=2.16, p<0.05), which was greater than that revealed in the saline session (t(28)=1.99, p=0.06). The analyses on the VStr revealed significant food value-related activity in both the ghrelin and saline conditions (p’s<0.05; n=29; see Figure 4C). Moreover, in the right VStr, the CS Value-associated activity was stronger during food versus non-food odor conditioning (t(28)=2.01, p=0.05, ghrelin; t(28)=1.87, p=0.07, saline). However, only in the ghrelin condition, food CS Value-associated activity correlated with in-scanner hedonic scores in both the right and left VStr (for both correlations, r=0.51, p=0.06; n=14; Figure 4D).

**Figure 4.**
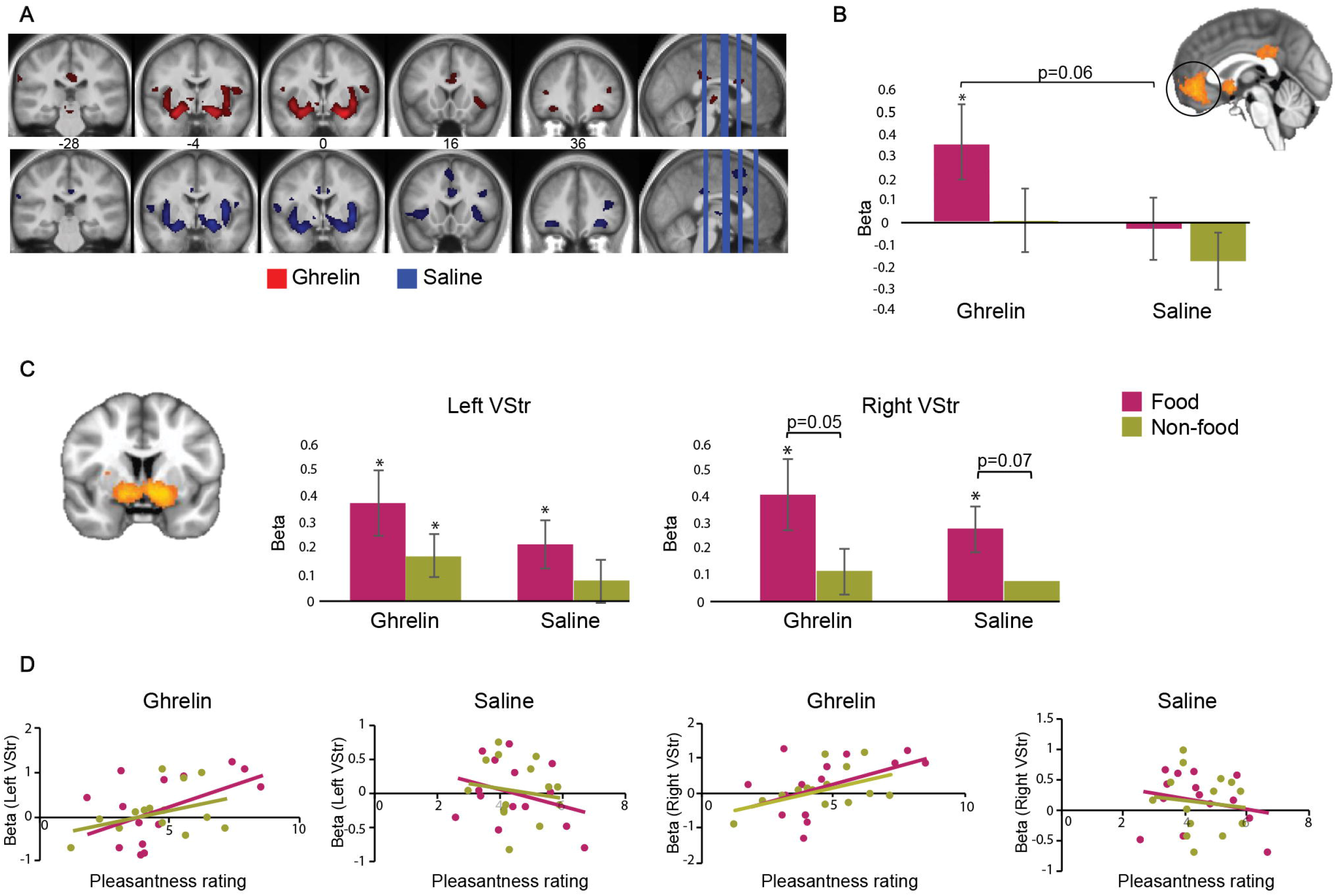
Ghrelin heightens brain response associated with expected value assigned to food cues. (A) Expected value assigned to cues (CS Value) correlated with activity in many brain regions including the piriform cortex, insula, anterior and posterior cingulate cortex and OFC in both ghrelin and saline conditions. (B) In the analysis focusing on the part of the vmPFC previously associated with subjective value, we observed greater CS Value-related activity during food conditioning in the ghrelin versus saline condition (t(28)=1.99, p=0.06). (C) Another analysis focused on the clusters previously identified to encode subjective value that largely include the VStr. We observed increased CS Value-related activity on food trials following ghrelin and saline administration in both the left and right VStr (p’s<0.05). (D) Only in the ghrelin condition, food CS Value-related activity in the VStr was correlated with in-scanner pleasantness ratings (r’s=0.51, p=0.06).

**Table 2.**
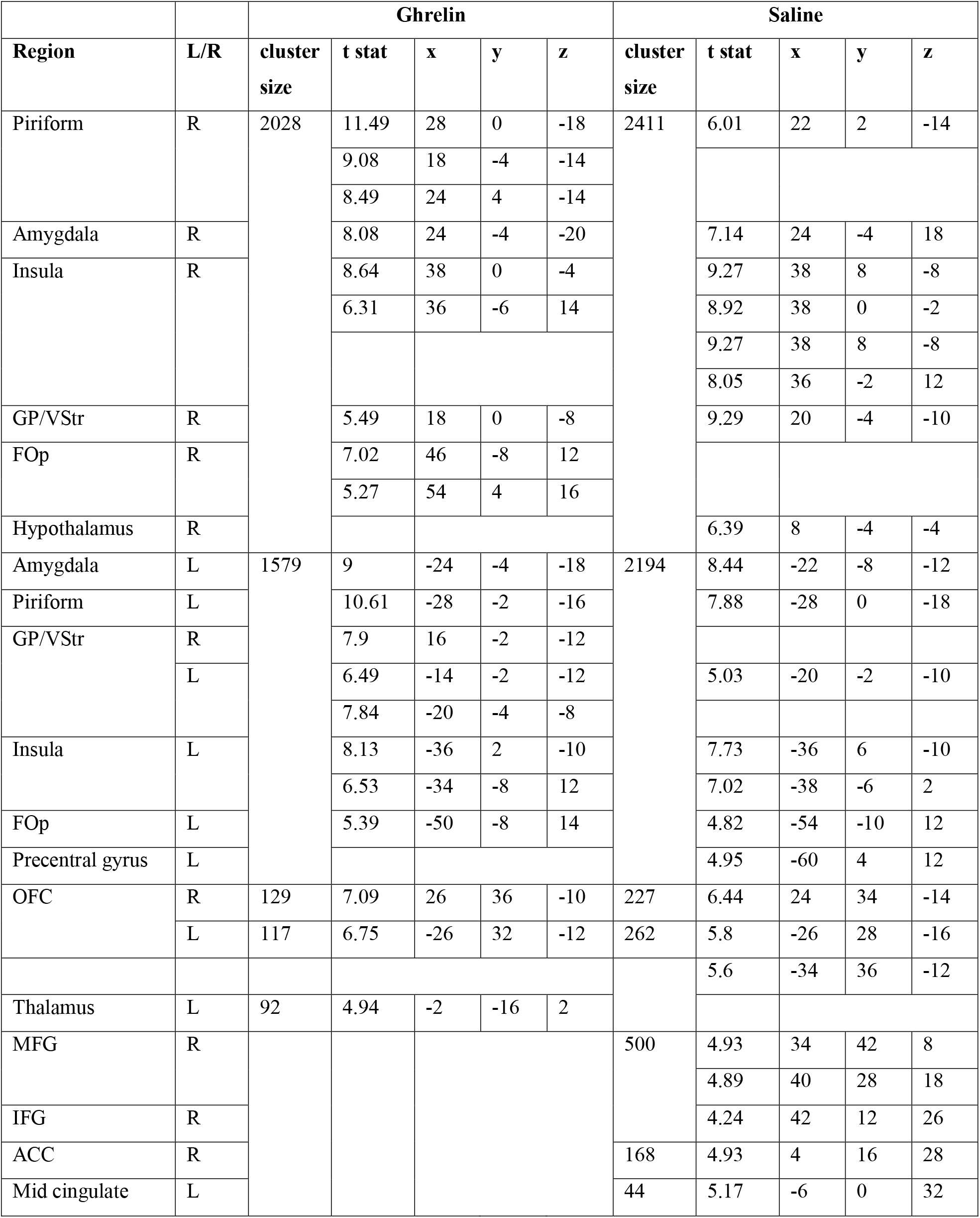
Value-associated activity. GP: globus pallidus, OFC: orbitofrontal cortex, MFG: middle frontal gyrus, IFG: inferior frontal gyrus, VStr: ventral striatum, ACC: anterior cingulate cortex, Fop: frontal operculum.

In line with these neuroimaging findings, the delayed rating task revealed that the abstract images paired with food odors following ghrelin injection were perceived to be more pleasant. This, considering our RT and RPE-related results, supports ghrelin-induced enhancement of conditioning to food odors, leading to increased incentive value of the conditioned stimuli paired with food odors.

### Ghrelin Strengthens Hippocampus-Ventral Striatum Coupling during Food Conditioning

Complex cognitive processes such as learning tend to recruit networks of spatially separate brain regions rather than engaging them independently. Indeed, connectivity between the hippocampus and VStr has been shown to support value-related learning by linking stored memories of value in the hippocampus to reinforcement processes in the striatum (Wimmer and Shohamy, 2012). We therefore conducted a generalized psychophysiological interaction (gPPI) analysis (McLaren et al., 2012) to see if ghrelin modulated task-dependent connectivity between the hippocampus and VStr regions that were revealed in the activation analysis to be associated with RPE. We observed, on trials where odors were delivered, a significantly greater coupling between the left VStr (seed) and the left hippocampus on food-odor trials in the ghrelin versus saline condition (t(28)=2.14, p=0.04; see Figure 5). In the saline condition, the left VStr was more strongly associated with the right hippocampus on non-food compared to food trials (t(28)=3.03, p=0.005).

**Figure 5.**
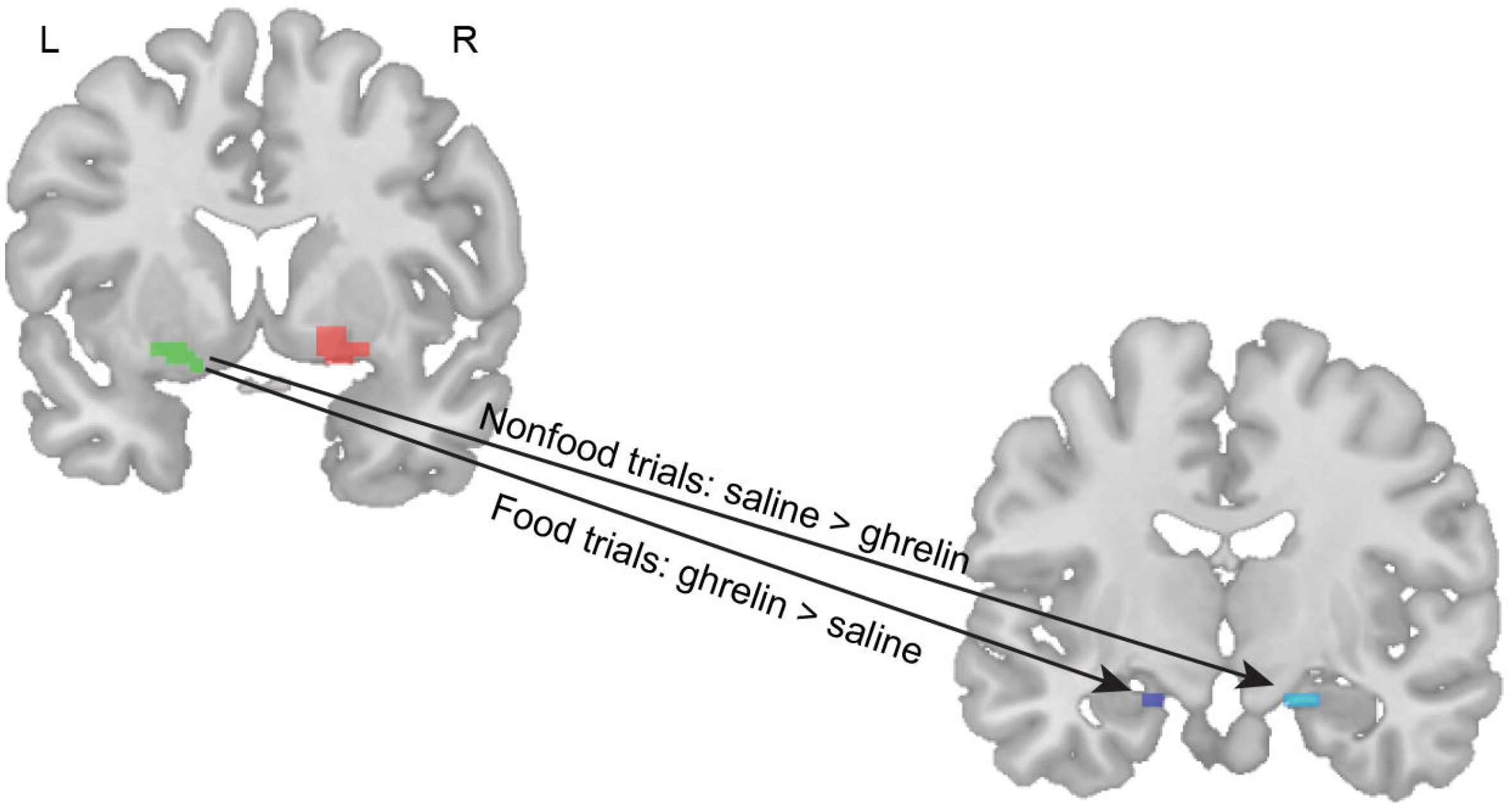
Ghrelin strengthens hippocampus-ventral striatum coupling during food conditioning. The gPPI analysis with the VStr as the seed region and the hippocampus as the target revealed a greater functional coupling between the left VStr and the left hippocampus during food conditioning in the ghrelin versus saline condition (t(28)=2.14, p=0.04). In the saline condition, the left VStr showed greater functional coupling with the right hippocampus on non-food compared to food trials (t(28)=3.03, p=0.005).

The above-mentioned brain regions thought to generate learning-related RPE signals are heavily connected to the hippocampus. The hippocampus is speculated to provide input into the VStr to modulate learning-related signals and participate in encoding and retrieving of cue-reward associations (Pennartz et al., 2011). For instance, in reinforcement learning tasks that rely on episodic memory for cues, associations have been found between learning performance and both stronger hippocampal activity and hippocampus-striatum coupling (Davidow et al., 2016; Wimmer and Shohamy, 2012). Ghrelin also acts on the hippocampus, where GHSR are densely expressed (Mani et al., 2014). Thus, ghrelin’s learning- and value-promoting effects also appear to be exerted via the hippocampus. Our study involved second order conditioning, which is thought to necessitate recruitment of the hippocampus (Wimmer and Shohamy, 2012). Indeed, we observed greater RPE-related activity in the hippocampus during food conditioning following ghrelin compared to saline injection. Furthermore, functional connectivity between the two regions associated with RPE, namely the hippocampus and VStr, was stronger on food versus non-food trials following ghrelin infusion while the reverse pattern was seen in the saline condition. Finally, the food trial-induced coupling between the two regions was significantly stronger following ghrelin compared to saline treatment.

The hippocampus is implicated in cue potentiated feeding, in which a food-paired conditioned stimulus drives feeding behavior (Kanoski et al., 2013). It is also necessary when contextual information must be used for the learning or expression of an association between a food cue and feeding behavior (Kanoski and Grill, 2017). Both phenomena depend on ghrelin signaling in the hippocampus. For example, ghrelin, as a meal anticipatory signal, has been shown to promote cue-driven feeding via actions on the hippocampus (Hsu et al., 2016), and GHSR blockade prevents cue-potentiated feeding (Walker et al., 2012). In animals trained on a fixed meal schedule, hippocampal GHSR blockade reduces food consumption at the anticipated mealtimes (Hsu et al., 2015), presumably by decoupling the temporal context from cue reactivity. There is also evidence from animal experiments that ghrelin acts during the formation of food-cue reward associations (Hsu et al., 2018; Walker et al., 2012). Thus, the hippocampus incorporates information about familiar food cues, the current context, and circadian and energy balance information to control feeding behavior. Animal studies implicate connections between hippocampus and mesolimbic dopamine structures including the VStr in these processes (Kanoski and Grill, 2017). Our results support this model, whereby ghrelin promotes the formation of context-specific cue-reward associations by augmenting hippocampal signaling and connectivity to VStr.

### The Actions of Ghrelin Are Food-specific

An intriguing finding revealed consistently across the dataset is that only the responses to food odors were modulated by ghrelin injection. As argued above, the effects on RPE and value appear to be plausibly exerted via DA signaling, which is known to be stimulated by ghrelin. However, given DA’s responsivity to a wide variety of rewards, it might be assumed that ghrelin’s actions could generalizable to non-food stimuli (Daniel and Pollmann, 2014). Indeed, a few studies have demonstrated ghrelin-induced modulation of responses to drug rewards such as cocaine and alcohol (Jerlhag et al., 2009, 2010; Wellman et al., 2005). In the present work, however, ghrelin injection enhanced learning with food, but not non-food, odors despite their similar pleasantness, intensity ratings and evoked brain responses. Ghrelin’s ability to selectively facilitate associative learning with food reward was revealed in reaction times, RPE and Value-related activity in dopaminergic brain regions, and in hippocampal-striatal connectivity. It is possible that ghrelin preferentially targets food-specific pathways within the DA system and other regions such as the lateral hypothalamus, which contains the highest density of GHSR and regulates appetite and energy balance (Olszewski et al., 2003; Toshinai et al., 2003). Lateral hypothalamic projections to VTA DA neurons (Korotkova et al., 2003; Nakamura et al., 2000) could then mediate this food-specific learning effect of ghrelin.

Alternatively, hippocampal involvement may also explain the food-specificity of our findings. Food odors are learned contextual cues that rely on hippocampal memory systems. Ghrelin may activate hippocampal memory traces of food-specific cues to promote associative learning via hippocampal-striatal connectivity (Kanoski and Grill, 2017). However, the precise neuronal mechanisms underlying ghrelin’s selective effects on food stimuli cannot be addressed here given the low spatial resolution of fMRI and our study design.

### Ghrelin Does Not Alter Odor Perception

There is some evidence that ghrelin can increase olfactory sensitivity and sniffing as it binds to GHSR present in the olfactory bulb and other odor-processing brain regions (Tong et al., 2011). In order to see if ghrelin’s effects on food-related learning are attributed to its influence on sensory signaling, neural activation associated with odor perception was examined by contrasting odor and air trials. Exposure to odors increased activity in the piriform cortex, insula, OFC, middle and inferior frontal gyri, VStr and posterior cingulate cortex in both ghrelin and saline conditions, which did not differ from each other (see Figure S4A, Table 3). Moreover, odor detection thresholds taken after scan did not differ between the ghrelin and saline conditions (t(17)=1.02, p=0.32). Finally, when fMRI response to different types of odors was investigated, food odors compared to air evoked activity in the piriform cortex, OFC, insula, ventral striatum, and middle frontal gyrus in both ghrelin and saline conditions (see Figure S4B, Table 4). Only following saline injection, non-food odors led to increased activation in the middle frontal gyrus. Taken together, we may conclude that ghrelin’s effects on food-related conditioning observed using our task cannot be attributed to increased sensory signaling.

**Table 3.**
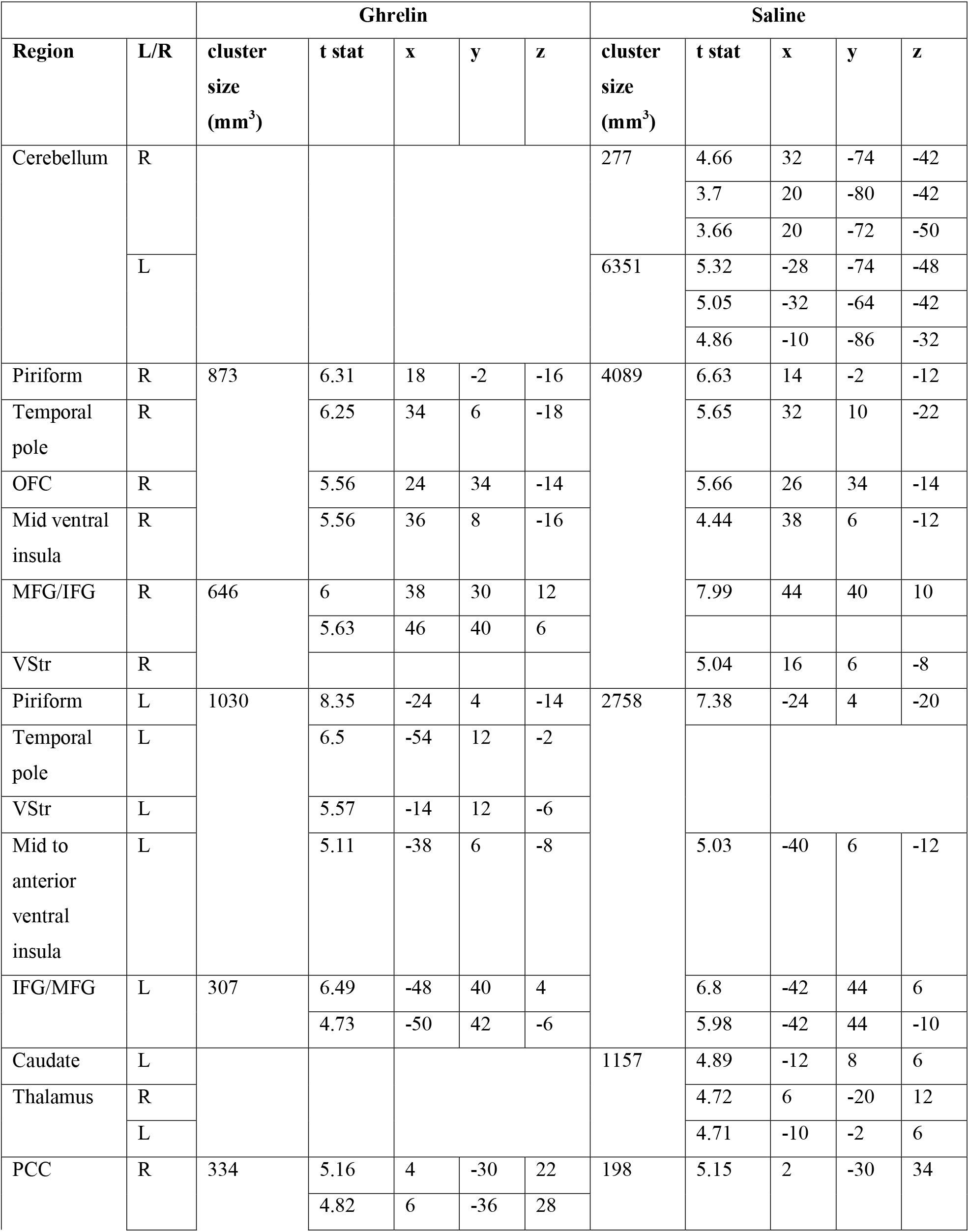
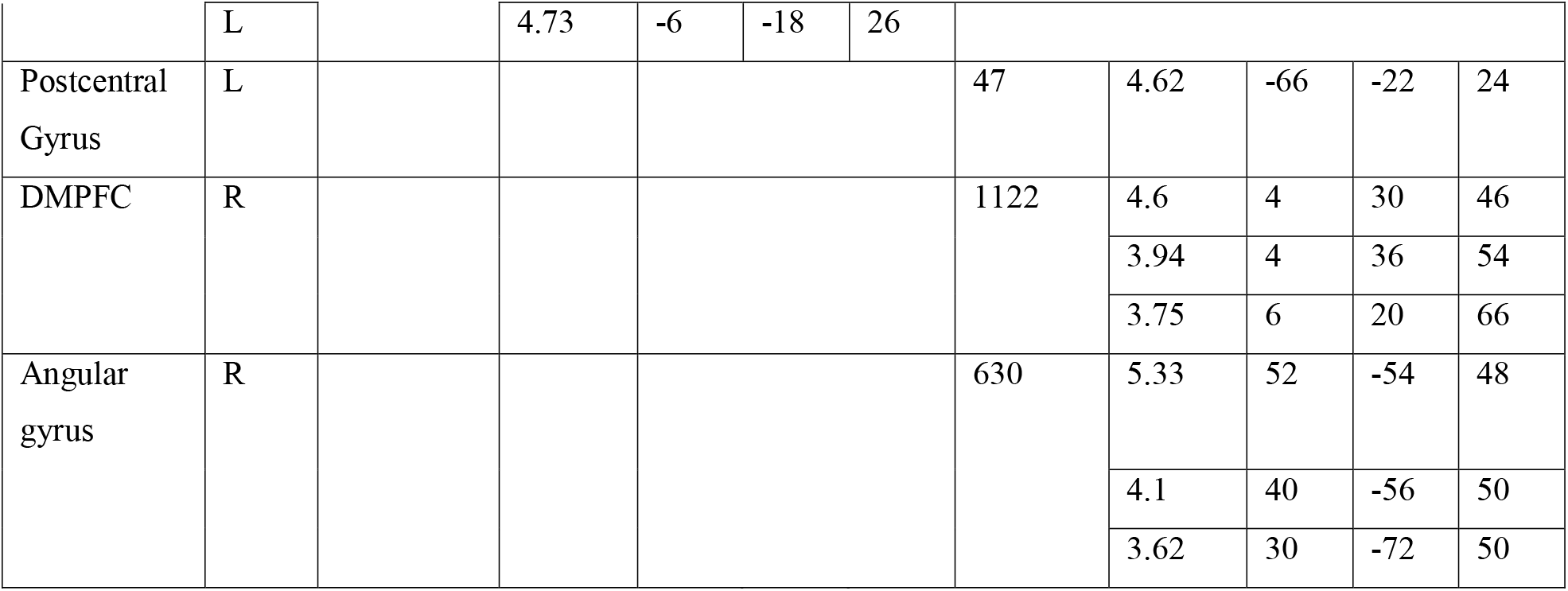
Odor-evoked brain responses. OFC: orbitofrontal cortex, MFG: middle frontal gyrus, IFG: inferior frontal gyrus, VStr: ventral striatum, PCC: posterior cingulate cortex, DMPFC: dorsomedial prefrontal cortex.

**Table 4.** BOLD responses to food and non-food odors. GP: globus pallidus, OFC: orbitofrontal cortex, MFG: middle frontal gyrus, IFG: inferior frontal gyrus.

|  |  |  | Ghrelin |  |  |  |  | Saline |  |  |  |  |
| --- | --- | --- | --- | --- | --- | --- | --- | --- | --- | --- | --- | --- |
|  | Region | L/R | cluster size | t stats | x | y | z | cluster size | t stats | x | y | z |
| **Food odors** | Ventral Insula | R | 80 | 5.52 | 34 | 6 | -18 |  |  |  |  |  |
|  | Piriform | R | 125 | 5.88 | 16 | -4 | -16 |  |  |  |  |  |
|  | GP | R |  |  | 26 | -6 | -8 |  |  |  |  |  |
|  | Piriform | L | 207 | 7.01 | -26 | 2 | -16 | 80 | 6.25 | -20 | 2 | -18 |
|  | Midbrain | L |  | 4.61 | -12 | -10 | -14 |  |  |  |  |  |
|  | GP | L |  | 4.28 | -14 | -4 | -8 |  |  |  |  |  |
|  | Temporal pole | L | 62 | 6.28 | -56 | 10 | -4 |  | 4.56 | -30 | 8 | -20 |
|  | OFC | R | 83 | 5.22 | 22 | 32 | -14 |  |  |  |  |  |
|  |  |  |  | 4.25 | 32 | 36 | -10 |  |  |  |  |  |
|  |  |  |  | 3.93 | 22 | 40 | -8 |  |  |  |  |  |
|  | IFG/MFG | R |  |  |  |  |  | 392 | 6.19 | 42 | 36 | 8 |
|  |  |  |  |  |  |  |  |  | 5.98 | 46 | 36 | 16 |
|  |  |  |  |  |  |  |  |  | 5.17 | 44 | 44 | 6 |
|  | Cerebellum | L |  |  |  |  |  | 41 | 5.36 | -38 | -60 | -24 |
| **Non-food odors** | MFG/IFG | R |  |  |  |  |  | 154 | 5.98 | 42 | 44 | 6 |
|  |  |  |  |  |  |  |  |  | 4.48 | 46 | 38 | 16 |

### Limitations

We propose that greater DA signaling explains accelerated food-related learning following ghrelin treatment. Our interpretation is based on substantial evidence that ghrelin stimulates the DA system that is thought to encode RPE-related activity associated with reinforcement learning. However, DA signaling was not directly assessed in this study. Moreover, there is some evidence that ghrelin modulates other systems such as the opioid and the endocannabinoid, which also interact with DA and may influence food motivation (Edwards and Abizaid, 2016; Kawahara et al., 2013). Therefore, the possibility for the involvement of non-dopaminergic systems in ghrelin-induced facilitation in learning should not be ruled out. On a related note, while ghrelin is known to stimulate release of other hormones such as GH, cortisol and adrenocorticotropic hormone, we attribute our findings to the effects of ghrelin, as pharmacological levels of the hormone were injected (Arvat et al., 2001). Nevertheless, readers should keep in mind the potential indirect influences of the other hormones on our results.

### Clinical Relevance

Our results support the animal literature in highlighting the role of ghrelin in the motivational and learned aspects of feeding. They may explain the consistent observation that, while chronic ghrelin administration causes weight gain, ghrelin or GHSR null mice are the same weight as wild-type animals when chow-fed (Müller et al., 2015). However, lack of ghrelin signaling appears to protect these animals against diet-induced obesity when they have access to appetizing high-fat foods. Ghrelin deficient mice may simply lack the ability to condition to high-calorie foods, despite having seemingly normal energy homeostasis.

Obesity is characterized by abnormal reactivity to food-related cues abundant in our environment (Boswell and Kober, 2016; Jansen et al., 2016; Stice and Yokum, 2016). Here we show that ghrelin enhances food-odor conditioning and its related BOLD response in mesolimbic projection sites. This provides a mechanistic link between energy signaling and learning about the food environment.

The ghrelin-responsive regions identified here have been implicated in a neural endophenotype that confers vulnerability to obesity. Cue reactivity in the vmPFC and VStr has been shown to encode the learned value of food cues based on their energy content (de Araujo et al., 2013; Tang et al., 2014) and this response in turns appears to correlate with obesity and prospective weight gain (Murdaugh et al., 2012; Stoeckel et al., 2008). In summary, conditioning to the hedonic, and typically caloric, aspects of food cues modifies the neural response to these cues in ways that appear to predispose to future weight gain. Our results show that homeostatic / circadian signals like ghrelin play a role in the neural plasticity processes that predispose to obesity. By providing further support for ghrelin’s role as a link between energy balance and motivation and learning, the present work unravels potential mechanisms through which ghrelin may contribute to both normal and maladaptive eating behaviours.

## STAR ⋆ METHODS

### Contact For Reagent And Resource Sharing

All statistical maps are available from the authors. Further information and requests for resources should be directed to and will be fulfilled by the Lead Contact, Alain Dagher.

### Experimental Model And Subject Details

#### Participants

Forty young healthy right-handed individuals (age: 22.46±2.60, body mass index: 23.33±2.98, 17 women) were recruited by advertisements. Of those, 38 completed the study. Exclusion criteria included psychiatric or neurological illness, body mass index > 25.9 (men) and >27.0 (women) or <19, gastrointestinal or eating disorders, current use of medications (other than oral contraceptives), tobacco or other drugs of abuse, food allergies, hay fever, deviated nasal septum, a cold or sinus infection, vegetarianism, and/or contraindications for MRI scanning. In order to exclude individuals with abnormal olfactory thresholds, we administered a brief olfactory test where participants were presented with 10 sets of three bottles (one with an odorant and the other two containing no odorant) and instructed to identify the bottle from each set that smells strongest. We also excluded individuals with abnormal eating behaviours who scored above 20 on the Eating Attitude Test (Garner and Garfinkel, 1979), and/or answered “Yes” to any of the two questions on the eating-related section of the Structured Clinical Interview for the Diagnostic and Statistical Manual of Mental Disorders-IV Screening Module (APA, 1998). Female participants were scanned during the luteal phase given the differential reward responses documented throughout the menstrual cycle (Dreher et al., 2007). All participants provided written informed consent as approved by the Montreal Neurological Institute (MNI) Research Ethics Board and received monetary compensation for their time and effort.

### Method Details

#### Ghrelin and task stimuli

Human ghrelin (C_149_H_249_N_47_O_42_, molecular weight = 3370.9) was obtained from Clinalfa (Bachem Distribution Services GmbH, Weil am Rhein, Germany). The hormone was manufactured according to GMP regulations and was sterile and pyrogen free. The peptide was delivered lyophilized in individual 100μg glass vials, and intended for intravenous injection to humans. Ghrelin was reconstituted with saline (1 ml).

Food odors (strawberries and cream, caramel, guava, and orange) and non-food odors (rose, olibanum, freesia, and muguet) used in the study were matched for intensity, familiarity and pleasantness based on a pilot study using 28 different commercially available odorants conducted in a separate group of 15 volunteers. Odors (25ml each, undiluted odorants) were delivered through a computer-controlled, 8-channel olfactometer (Dancer Designs, Merseyside UK), which ensures accurate odor onset and a steep odor rise-time. The visual stimulus set, taken from the Abstract Design List learning task (Jones-Gotman, 1986), consisted of 12 abstract line drawings, 6 made of straight lines and 6 of curved lines.

#### Testing Sessions

Each participant underwent two fMRI sessions following saline or ghrelin injection, scheduled at least one week apart at the same hour of the day. The order of ghrelin and saline injection was counterbalanced. Participants received saline or 1μg/kg of ghrelin intravenously, in single-blinded fashion. No side effects were reported.

As illustrated in Figure 1A, on testing day, participants arrived at the laboratory between 7:30AM and 11AM and were provided with a standard breakfast following a 12-hour overnight fast. The breakfast menu was designed to be moderately low in glycemic index and protein, to minimize their influence on brain function. The meal included 2 slices of toasted bread (1 white and 1 whole wheat), 42g of cheddar cheese, 10ml of butter, 125ml of orange juice and 1 cup of coffee or tea with 20ml of 2% milk and 1 sachet of white sugar. Participants were instructed to consume the provided meal in its entirety and nothing else until the end of the session. Immediately before and after breakfast, subjects were asked to rate their levels of hunger, boredom and irritability on a visual analog scale (VAS), ranging from −5 (not at all) to 5 (extremely).

Neuroimaging took place 3 hours after the breakfast when the circulating ghrelin levels are expected to be at nadir (Cummings et al., 2001). Prior to scanning, subjects completed the Profile of Mood States questionnaire (McNair, D.M. et al., 1971) and again reported their hunger, boredom and irritability levels. Subsequently, we collected participants’ saliva and blood samples in order to measure levels of cortisol (saliva), and insulin, growth hormone and glucose (blood). Ghrelin or saline was then administered by infusion into the antecubital vein over 60 seconds, after which another saliva sample and the VAS ratings (hunger, boredom and irritability) were collected. Participants were then placed in the MRI scanner. The session began with a 5-minute structural scan, followed by seven functional scans (7 minutes each) during which subjects performed the odor conditioning task detailed below.

Upon completion of the imaging part of the study, we again administered the VAS scales to quantify participants’ hunger, irritability and boredom and collected their saliva and blood samples. In a subset of participants (n=18), we also assessed odor detection thresholds for n-Butanol (Fisher Scientific Pittsburgh, PA) using a staircase, triple-forced choice procedure as described by Kobal and colleauges (Kobal et al., 2000).

After approximately 24 hours following the scan session, participants returned to the laboratory for a behavioural session where they provided pleasantness and familiarity ratings for the conditioning images and two novel images, and pleasantness, familiarity and intensity ratings for the odors used during the scan as well as two new odors.

The fMRI and behavioural sessions took place 7 to 30 days apart. Participants completed the same tasks with different sets of visual and olfactory stimuli.

#### Blood and cortisol sampling

Blood samples were collected from the antecubital vein (approximately 2ml) before injection of ghrelin or saline and after the scan in order to quantify the serum levels of growth hormone, glucose and insulin. Blood was collected in gold-top serum separation tubes (bd.com) and placed on ice immediately. Tubes were then sent to the McGill University Health Centre biochemical laboratory for analysis.

Salivary cortisol was sampled using the salivette collection device (Sarstedt Inc., Quebec City, QC, Canada) at three different time points, before and after saline or ghrelin injection and after the scan. Participants were required to place the salivettes in their mouths for approximately one minute. The samples were stored at −20C until analysis. Cortisol (nmol/l) was quantified using a time-resolved fluorescence immunoassay as described by Dressendorfer and colleagues (Dressendörfer et al., 1992).

#### fMRI olfactory conditioning task

The fMRI task design was based on Gottfried et al. (Gottfried et al., 2002a, 2002b). Four odors (2 food, 2 non-food) were paired with 4 abstract images (CS+) on a 50% positive reinforcement schedule. The remaining two images (CS-) were paired with odorless air. Stimuli, their pairings and the presentation sequence varied between the two sessions and across the participants. Stimuli were presented in a pseudo-random order such that no two identical images or odors appeared consecutively. In addition, no more than five air-paired events were presented in a row.

As illustrated in Figure 1B, each trial began with the 1250ms-presentation of a visual stimulus. Its corresponding odor was delivered 500ms after the image onset and disappeared together with the image. Each trial was followed by an inter-trial blank screen with a jittered interval ranging from 6500ms to 8500ms. Upon viewing each image, subjects indicated whether the image was made of curvy or straight lines using a MRI-compatible mouse-like device. There were 7 fMRI runs in each session, each of which was composed of 36 trials (12 CS+paired, 12 CS+ unpaired, &12 CS-). At the end of each functional run, a subset of participants (n=21) rated pleasantness of the 6 images on a Likert scale, ranging from 0 (not pleasant at all) to 10 (highly pleasant).

#### fMRI data acquisition

Imaging data were acquired using a 3T Siemens (Erlangen, Germany) Magnetom Trio MRI scanner with a 32-channel head coil. Following a MPRAGE, T1-weighted anatomical scan (Voxel size = 1×1×1 mm), functional T2*-weighted echoplanar images were acquired using blood oxygenation level dependent (BOLD) contrast (7 sessions of 140 volumes each, 38 axial slices, TR = 2300ms, TE = 30ms, Flip angle = 90°, Voxel size = 3.5×3.5×3.5ms, FoV = 224mm). E-Prime (Psychology Software Tools, Pittsburgh, PA) running on a PC laptop was used to trigger the olfactometer and to present visual stimuli, projected onto a screen in the fMRI scanner visible to subjects through a mirror system, and to record subjects’ button responses

### Quantification And Statistical Analysis

#### Modeling of reward prediction error signals

We used the Rescorla-Wagner reinforcement learning model to generate trial-by-trial prediction signals, namely expected value assigned to CS (referred to as “CS Value”) and RPE (Rescorla, R.A. and Wagner, A.R.). The RPE signal, δ, is defined as the difference between the value of the actual outcome on a given trial, R, at time t, and that of the expected outcome on that trial, V.

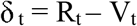

We modeled the presentation of an odor with R=10, and the omission of an odor with R=0, for both food and non-food odors. The CS Value (V) was updated by adding δ weighted by a learning rate α: V_t_ = V_(t-1)_ + α δ_t_. We derived α from participants’ reaction times (RT) that were used as trial-by-trial measure of conditioning (Wilson and Niv, 2015). It has been previously shown that learning systematically modulates RT. Changes in RT also reflect changes in the coded value of each stimulus, and consequently values estimated by reinforcement learning models (Critchley et al., 2002; Gottfried, 2003; Gottfried et al., 2002a; Seymour et al., 2004). Based on the assumption that the RT would vary as a function of the current CS Value, we estimated trial-by-trial values for a range of learning rates (0 to 1) and regressed RT onto each of the value curves (O’Doherty et al., 2007). The learning rate that minimized the error was 0.17.

#### Behavioural analysis

Participant RTs and pleasantness ratings were used as indices of learning (Critchley et al., 2002; Gottfried et al., 2002a; Howard et al., 2015). We speculated that the images associated with food odors following ghrelin injection would induce faster RTs and greater pleasantness. RTs were z-score normalized, after which odor type-specific RTs for each participant were averaged for each conditioning run. A three-way ANOVA was conducted to observe the effects of odor type, time and treatment on RT using SPSS (version 23, SPSS Inc., Chicago, IL). RT data was analyzed in 29 participants whose fMRI dataset was deemed valid (see below).

Owing to a size-related error in the images presented during the hedonic rating tasks administered to seven participants, the hedonic rating data were analyzed in 14 participants who were shown properly-sized images. Event type-specific in-scanner hedonic ratings for each subject were averaged for each conditioning run, and were then analyzed using a three-way ANOVA. Additionally, we performed a two-way ANOVA on the pleasantness rating data collected following a 24-hour delay to investigate the effects of event type and treatment.

#### fMRI data analysis

Neuroimaging analyses were conducted in 29 participants as nine were excluded due to missing responses during the conditioning task and/or excessive head movements (n=8), and lack of growth hormone response to ghrelin injection (n=1).

SPM 8 software (Wellcome Department of Imaging Neuroscience, London, UK) was used for preprocessing and statistical analysis of the fMRI data. The images were slice-time corrected, realigned to the first volume, and normalized into MNI space (final voxel size = 2 × 2 × 2 mm) (Evans et al., 1994). Spatial smoothing (isotropic Gaussian kernel of 6mm FWHM) was then performed to improve the signal-to-noise ratio of the images. Low frequency temporal drifts were removed using a high pass filter with a cut-off of 1/128s. The event-related general linear model (GLM) implemented by SPM was used for statistical analysis.

The first analysis was conducted to examine brain activity related to odor processing. We defined five event types: (1) air, (2) images paired with air, (3) odors (both food and non-food), (4) images paired with odors, and (5) button press. To investigate BOLD response during processing of different types of odors, we built another model with the following event types: (1) air, (2) images paired with air, (3) food odors, (4) images paired with food odors, (5) non-food odors, (6) images paired with non-food odors, and (7) button press.

Several parametric analyses were additionally conducted to examine CS Value- and RPE-associated brain activity. First, we defined three event types: (1) images, (2) odors (including odorless air), and (3) button press. In order to identify brain areas whose fMRI activity is modulated by stimulus values regardless of the type of odor, we entered CS Values (estimated by the reinforcement learning model) as parametric regressors for each trial at the time of the presentation of the image (Büchel et al., 1998). In another GLM, RPE signals were entered as parametric regressors for each trial at the time of the delivery of the odor.

With an aim to observe CS Value- and RPE-related brain activity in different conditions, we defined the following event types: food odor-paired abstract images (CS+), food odors, non-food odor-paired abstract images (CS+), non-food odors, air-paired images (CS-) never paired with an odor, air, and button press. In one GLM, CS Value was entered as a parametric regressor for each conditioning trial at the time of the presentation of an image. In the second GLM, RPE was entered as a parametric regressor for each corresponding trial at the time of the delivery of the odor or air.

For each of the analyses mentioned above, regressors of interest for the BOLD response were generated by convolving the modulated stimulus functions with a standard synthetic hemodynamic response function. The single-subject models also included the six movement parameters obtained from the realignment procedure. For each participant, linear contrasts of parameter estimates for conditions of interest were generated and subsequently submitted to a whole-brain second-level random effects analysis.

Additionally, we conducted region of interest (ROI) analyses on regions previously identified to be associated with subjective value (Bartra et al., 2013) and RPE (Chase et al., 2015), based on published meta-analyses. The analyses were performed using the MarsBaR toolbox (http://marsbar.sourceforge.net/). We obtained, for each session and for each participant, effect sizes for the contrasts of interest for each ROI, which were further analyzed using one-sample t-tests and paired t-tests in SPSS. An additional ROI analysis with small volume correction (SVC) was performed on the hippocampal regions defined by the AAL atlas implemented in SPM8 (Tzourio-Mazoyer et al., 2002).

Furthermore, to test if ghrelin modulated task-dependent connectivity between learning-related brain regions, we used a generalized form of psychophysiological interaction analysis (gPPI) (McLaren et al., 2012). As per our hypothesis, the regions of interest chosen for this analysis were activation clusters within the hippocampus and ventral striatum (VStr) that exhibited a significant modulation by overall RPE in both ghrelin and saline conditions at the group level (FDR corrected p<0.05). First, the physiological variable was derived from extracting de-convolved time series from the VStr seed for each subject. The psychological regressors were created by convolving the canonical hemodynamic response function with the onset times for food odor-paired images, food odor-unpaired images, nonfood odor-paired images, non-food odor-unpiared images, air-paired images, and button press. Subsequently the time series from the psychological regressor were multiplied with the physiological regressor, creating the interaction terms (PPIs). We were interested in functional connectivity between the two regions revealed in our activation analysis to be associated with RPE. Therefore, we took a ROI approach where the mean contrast estimates of the PPI regressor were extracted from the target ROI, namely the hippocampus. Repeated ANOVAs and paired t tests were then conducted on the contrast estimates.

## Acknowledgment

We thank Mercedes Pilkington for her assistance with collecting pilot study data and Dr. Donald McLaren for his help with the gPPI analysis. This study was funded by an operating grant from the Canadian Institutes for Health Research to A.D.

## Author contributions

S.M., J.B., A.D., T.M., and M.J. designed the study. J.H., J.B., T.M., and J.F. conducted the experiment; J.H., K.L., and Y.Z. analyzed the data; J.H. and A.D. wrote the paper.

## Declaration of interests

The authors declare no competing interests.

